# Expert Curation of the Human and Mouse Olfactory Receptor Gene Repertoires Identifies Conserved Coding Regions Split Across Two Exons

**DOI:** 10.1101/774612

**Authors:** If H. A. Barnes, Ximena Ibarra-Soria, Stephen Fitzgerald, Jose M. Gonzalez, Claire Davidson, Matthew P. Hardy, Deepa Manthravadi, Laura Van Gerven, Mark Jorissen, Zhen Zeng, Mona Khan, Peter Mombaerts, Jennifer Harrow, Darren W. Logan, Adam Frankish

## Abstract

Olfactory receptor (OR) genes are the largest multi-gene family in the mammalian genome, with over 850 in human and nearly 1500 genes in mouse. The expansion of the OR gene repertoire has occurred through numerous duplication events followed by diversification, resulting in a large number of highly similar paralogous genes. These characteristics have made the annotation of the complete OR gene repertoire a complex task. Most OR genes have been predicted *in silico* and are typically annotated as intronless coding sequences. Here we have developed an expert curation pipeline to analyse and annotate every OR gene in the human and mouse reference genomes. By combining evidence from structural features, evolutionary conservation and experimental data, we have unified the annotation of these gene families, and have systematically determined the protein-coding potential of each locus. We have defined the non-coding regions of many OR genes, enabling us to generate full-length transcript models. We found that 13 human and 41 mouse OR loci have coding sequences that are split across two exons. These split OR genes are conserved across mammals, and are expressed at the same level as protein-coding OR genes with an intronless coding region. Our findings challenge the long-standing and widespread notion that the coding region of a vertebrate OR gene is contained within a single exon.

## INTRODUCTION

Olfactory receptor (OR) genes represent 2% and 5% of the total number of protein-coding genes in human and mouse respectively, comprising the largest multi-gene family in mammalian genomes. ORs are G-protein-coupled receptors expressed by olfactory sensory neurons (OSNs) located in the olfactory epithelium in the nasal cavity, and bind to odorants (1). Each mature OSN expresses only one OR gene (2), leading to a diverse population of OSNs, each characterised by the specific OR protein they express. The olfactory system is tasked with the detection of an immense number of odorants with widely varying structures, and has evolved a diverse repertoire of OR genes to do so. OR gene expansion has been the result of numerous duplication events, generating clusters of paralogous genes that are often very similar to each other (3, 4). This OR gene expansion resulted in high frequencies of recombination, translocation, and gene conversion events. However, OR genes from different subfamilies can substantially differ in their protein sequence, with similarities as low as 35% (5). Annotation of the OR gene repertoire has therefore been a complex task. Determining orthologous and paralogous relationships often requires careful consideration of the sequence identity between closely related proteins. Furthermore, species-specific expansions of particular OR clades are common (6, 7), and even within the same species there is genotypic and haplotypic variation in the encoded OR repertoire across individuals of a population (8–10).

Currently there are numerous disparities between databases as to whether an OR locus is protein-coding or pseudogenised, as well as on the length of the coding sequence. Historically, OR coding sequences have been described as intronless and until recently, most OR genes were annotated as single-exon structures. However, transcriptomic evidence from RNAseq studies of the olfactory mucosa of several mammals has revealed that OR genes have complex gene structures, with multiple exons and widespread alternative splicing (11–13). In this study, we present the outcome of an extensive expert annotation effort, to comprehensively characterise the human and mouse OR gene repertoires, adding previously missed genes and amending the protein-coding or pseudogene status of many loci. Additionally, we used RNAseq data to build gene models for a large fraction of the repertoires of both species. Most importantly, we identified 13 human and 41 mouse OR genes that contain an intact coding sequence split across two exons, a number of which were previously thought to be pseudogenes.

## MATERIALS AND METHODS

### Olfactory receptor gene annotation pipeline

The expert OR gene annotation pipeline is summarised in Figure 1. Manual annotation was performed in our in-house Otter annotation system (Genome Research Ltd. https://www.sanger.ac.uk/science/tools/otter), which employs the ZMap graphical user interface (14, 15). We started by retrieving all annotated OR genes in the human and mouse genomes from RefSeq (16) (https://www.ncbi.nlm.nih.gov/refseq/), MGI (17) (http://www.informatics.jax.org/), and HORDE (18) (https://genome.weizmann.ac.il/horde/), a database of human OR gene sequences (19). Most OR genes are arranged in clusters along the genome, which are prone to duplication and recombination events that lead to the expansion of the repertoire. To identify any missing OR loci, we first extracted all genomic sequences with an annotated OR gene or gene cluster, along with the flanking 500 kb. We then used dotter (20) to compute the alignment of a few representative OR genes from each extended genomic region. These were visualised as dot matrix plots and allowed the identification of any matches that were not already annotated as OR genes. These putative novel OR loci were included in the corresponding OR gene repertoires for downstream analyses.

**Figure 1.**
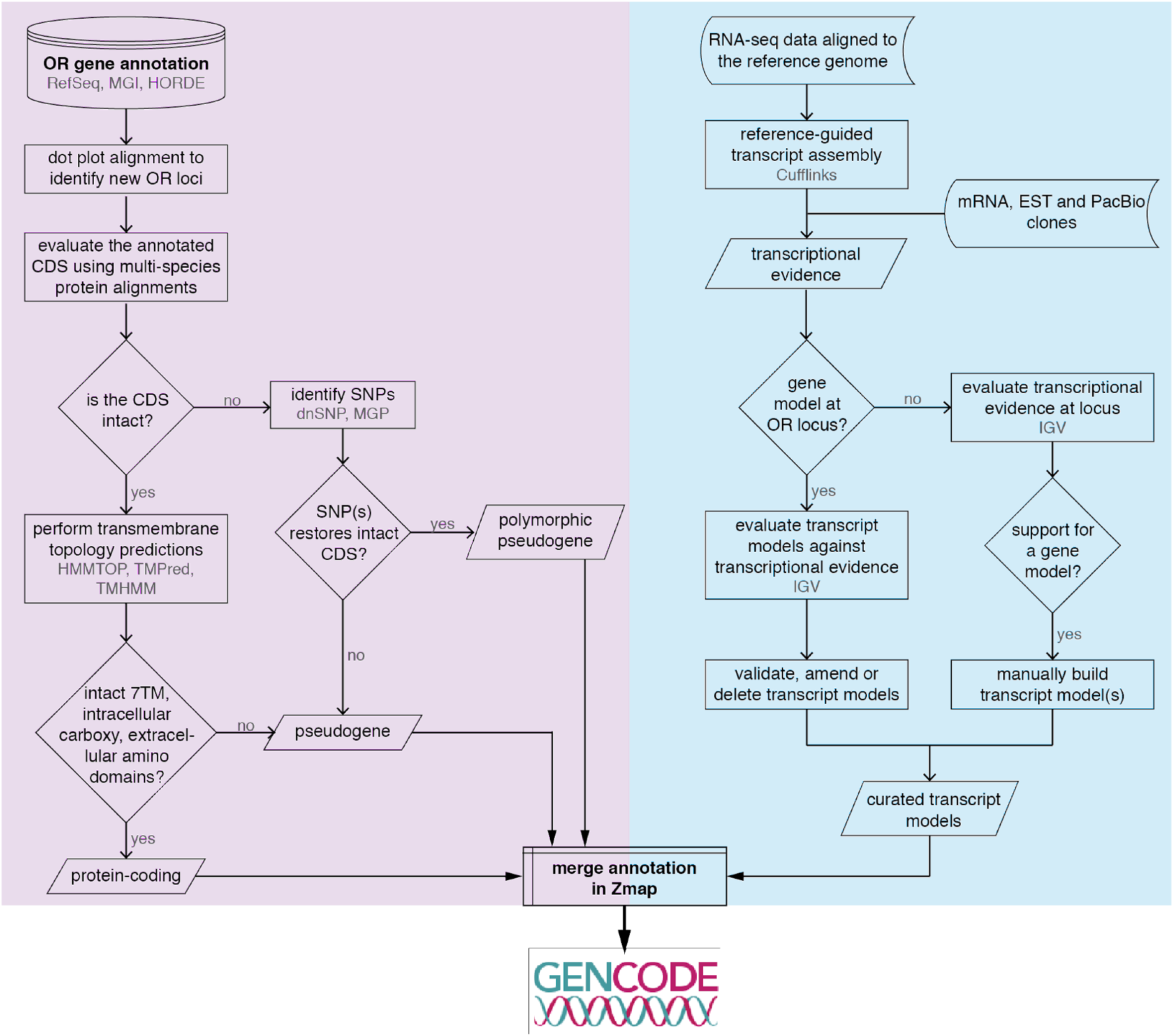
Olfactory Receptor annotation pipeline. Flow diagram showing the steps taken to annotate all OR loci of the human and mouse genomes. Specific databases and programs used are indicated in grey. The pipeline consists of two major tasks: 1) curating all available annotation for OR genes, as depicted on the purple-shaded steps; and 2) integrating transcriptional evidence from RNA-seq data and mRNA, EST and PacBio clones to construct gene models including untranslated regions, as shown on the blue-shaded steps. Results from both tasks were integrated into a comprehensive annotation of the human and mouse OR repertoires. These were subsequently added to the GENCODE project. 7TM = seven transmembrane domain.

For each OR locus, we performed cross-species conservation analyses, including human, mouse, cow, sheep, cat, donkey, shrew, mole, guinea pig, elephant, dog, sheep, rat, chimp, gorilla, orangutan, marmoset, lemur, bushbaby, tarsier, baboon, bonobo and/or gibbon. These analyses served to determine the biotype (protein-coding or pseudogene) and, if necessary, amend the length of the coding sequence (CDS) and/or splice sites sourced from the databases. Specifically, unambiguous orthologues were identified by performing a BLAT search (21) of the protein sequence of each OR gene in the UCSC browser (22); if none were found, the closest paralogue(s) was used instead. When alternative open reading frames (ORFs) were present, we favoured the initiation codon (iATG) with highest conservation. An alternative protein isoform for a locus was created if it had a predicted intact ORF, conserved OR topology and high identity to known annotated isoforms.

OR loci that contained an intact ORF were then subjected to transmembrane topology predictions using HMMTOP (23), TMPred (24) and TMHMM Server v 2.0 (25). OR genes with an intact ORF of at least 300 aa, with a predicted seven-transmembrane domain structure (characteristic of OR proteins), an extracellular N-terminus and intracellular C-terminus were biotyped as protein coding.

For loci without an intact ORF, those with three or fewer disruptions were checked against NCBI dbSNP database (26) and The Mouse Genomes Project catalogue of mouse strains variation (27), to assess whether there were reported single nucleotide polymorphisms (SNPs) that restored an intact, full-length ORF. Restored loci were annotated as polymorphic pseudogenes. The remaining disrupted loci were designated as unprocessed pseudogenes. For the human pseudogenes, where a syntenic one-to-one protein-coding orthologue could be unequivocally established in the mouse (GRCm38) or dog (Broad CanFam3.1) genomes, the locus was annotated as a unitary pseudogene instead. We did not perform this analysis for the mouse repertoire since the large number of very closely related paralogues makes it difficult to establish unequivocal orthologous relationships. For all pseudogenes, the *pseudogenic CDS* was determined using protein homology with existing protein-coding ORs. This was achieved using multiple alignments that were visualised in the Blixem browser (28), with the genomic coordinates for homologous regions to protein-coding OR genes manually determined.

The second part of the annotation pipeline involved the use of RNAseq data from human and mouse olfactory mucosa samples to assemble mapped reads into transcript models using Cufflinks (see next section for details). Each annotated OR locus was visualised in the Integrative Genomics Viewer (IGV) (29, 30) guided by Ensembl and/or HORDE annotation, along with the Cufflinks gene models and RNAseq data from all samples combined together. Evidence from any available mRNA and/or EST clones from GenBank (irrespective of their tissue of origin) was also considered, where the alignment was best-in-genome. Each Cufflinks model was manually assessed, revealing numerous inaccuracies which were corrected using bespoke in-house software (https://gitlab.com/olfr/olfr_transcript_model_curation). Briefly, short (1-3 bp) read overhangs at splice junctions incorrectly generated non-canonical splice sites; spliced reads were often misaligned, with a short portion of the read mapping to a close paralogue instead of the target gene, resulting in erroneous chimeric transcript models; the 3’ and 5’ termini predictions were often extended too far, and thus UTRs were always terminated when coverage dropped below 3 reads; and, finally, failure to account for drops in coverage due to low complexity regions (often found within OR gene UTRs (13) resulted in premature termination of the transcript model.

For highly expressed loci, only the most abundant transcript models were retained. For a fraction of OR genes, we observed a drop in the average read depth towards the end of the 3’ UTR, suggesting the possibility of alternative 3’ termini. In these cases we used the longest UTR in our transcript models. We also took a conservative approach to define the 3’ UTRs of genes in close proximity on opposite strands, often terminating the transcripts where read depth differences occurred between the two loci.

Where no Cufflinks models were predicted for OR loci, we checked whether there was enough evidence from the RNAseq data to manually build transcript models. To build a transcript model we required contiguous overlapping reads (except in low complexity regions), support from a minimum of 3 RNAseq reads spanning the splice junctions, and defined the UTR ends when read depth dropped below 3 reads.

All transcript models were integrated into Zmap (14, 15) using Annotrack (31) and constitute our final annotation of the complete human and mouse OR gene repertoires. We refer to this annotation as Ensembl-HAVANA, which was then integrated into the Ensembl/GENCODE reference gene set (32, 33) that is available through the Ensembl and UCSC genome browsers. As both Ensembl and UCSC genome browsers import information from other datasets (e.g., RefSeq), some genes contain additional transcript models not included in the curated dataset from this paper (e.g., *Olfr240-ps1*, discussed later in this paper). We have provided both a detailed table with every annotated transcript in both species (Supplementary File 1) and GTF annotation files including only the Ensembl-HAVANA curated human (Supplementary File 2) and mouse (Supplementary File 3) OR gene models.

### RNAseq datasets

For mouse, we used previously published data from two studies (11, 34) comprising RNAseq of whole olfactory mucosa samples of adult (8-10 weeks old) male and female C57BL/6J mice (12 samples in total, 6 from each sex). Raw data were retrieved from the European Nucleotide Archive (ENA; https://www.ebi.ac.uk/ena) from projects PRJEB1365 (11) and PRJEB5984 (samples ERS658588, ERS658589, ERS658590, ERS658591, ERS658592 and ERS658593) (34).

For human, we used data from two published studies comprising RNAseq of olfactory mucosa biopsy samples (three male samples from (35); two male and two female samples from (12)). Raw data were retrieved from the European Genome-Phenome Archive (EGA, https://ega-archive.org/) study EGAS00001001486 (35) and from NCBI Gene Expression Omnibus (https://www.ncbi.nlm.nih.gov/geo/) project GSE80249 (12).

An additional six human olfactory mucosa samples were collected and sequenced to increase the coverage of the human repertoire, as previously described (35). Briefly, samples were collected from male patients undergoing endoscopic sinus surgery in Leuven, Belgium, for resection of an adenocarcinoma. During the procedure, olfactory mucosa of the contralateral (healthy) side was harvested from the olfactory groove. Collected tissue was stored in RNAlater and shipped to the Max Planck Research Unit of Neurogenetics (Frankfurt, Germany) for further processing. For one patient the sample was divided into three samples (samples 10-12), and each was processed separately. Thus, there was a total of six samples from four different patients (Supplementary File 6). All patients provided written informed consent according to the study protocol, approved by the Medical Ethical Committee on Clinical Investigations at the University Hospitals of Leuven on 23 April 2014 (S5648).

RNA was extracted using the RNeasy mini kit (Qiagen) following the manufacturer’s instructions. Then, mRNA was prepared for sequencing using the TruSeq RNA sample preparation kit (Illumina) with a selected fragment size of 200-500 bp. Samples were multiplexed together and sequenced on an Illumina HiSeq 2500 to produce paired-end 100 bp sequencing fragments. All raw data have been deposited in the EGA under accession EGAS00001001486.

### RNAseq data processing and analysis

Human and mouse RNAseq data were aligned to the corresponding reference genomes (GRCh38 for human and GRCm38 for mouse) using Tophat version 2.0.13 (36), with default parameters. Mapped reads were used to perform reference-guided transcript assembly with Cufflinks version 2.2.1 (37) with default parameters, guided by Ensembl annotation, version 85 for human and version 83 for mouse (38). For the mouse data, Cufflinks was run on every sample and the results were compiled into a unique set of gene models using Cuffmerge. For the human data, we increased the coverage of OR genes by merging all 9 samples from Saraiva et al. (2019) (35) and this study into one BAM file. Similarly, the four samples from Olender et al. (2016) (12) were merged into a second BAM file. Each of these files were then used as input for Cufflinks. The two sets of gene models were kept separate and curated in parallel.

Gene expression levels for the mouse repertoire were taken from (34) Figure 2 - source data 1 (https://doi.org/10.7554/eLife.21476.006). For the human repertoire we applied the same methods as in (34). Briefly, we obtained the number of fragments mapped to each gene using the script htseq-count (mode intersection-nonempty; HTSeq version 0.6.2; (39)), using Ensembl annotation version 95, which contains the Ensembl-HAVANA curated OR models (40). To account for differences in sequencing depth between samples, raw counts were normalised using the method implemented in the DESeq2 package (41). Further normalisation to account for differences in the proportion of OSNs per sample was performed using the method proposed by Khan et al. (2013) (42). This consists of using the geometric mean of five genes expressed specifically in OSNs *(ADCY3, ANO2, OMP, CNGA2* and *GNAL)* to compute a size factor to scale the normalised counts of OR genes. The code used to analyse the data can be found in https://github.com/xibarrasoria/ORgeneAnnotation_HAVANA. The normalised counts for the human OR repertoire are provided in Supplementary File 6.

**Figure 2.**
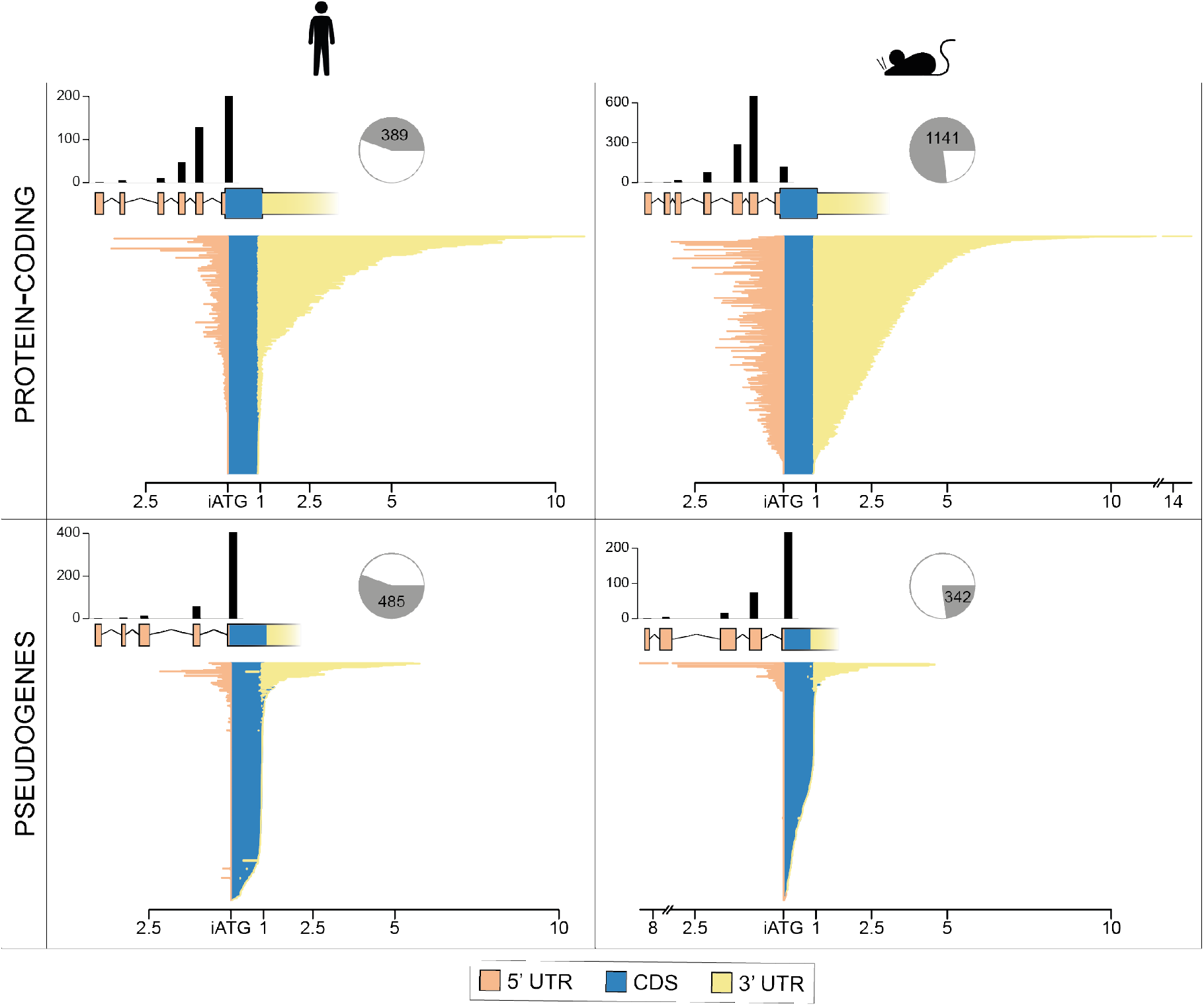
Structure and length of OR gene features. Barplots of the longest transcript for each OR gene, split by 5’ untranslated region (UTR), coding sequence (CDS) and 3’ UTR, in kilobases. Genes have been split into protein-coding (top) and pseudogenes (bottom), and by species (human on the right, mouse on the left). For pseudogenes, the CDS region of the barplot corresponds to the ***pseudogenic CDS.*** Above the barplot, a representative schematic of the OR gene structure; exons are shown as boxes and introns as connecting lines. Above the exons, bars indicate the number of genes with the corresponding number of exons; single-exon transcripts are rightmost, containing the CDS, and increasing number of exons progress to the left. The pie chart indicates the proportion of genes that are protein-coding or pseudogenised. iATG = initiation methionine of the open reading frame.

Data processing, statistical analyses and plotting were performed in R (43) (http://www.R-project.org).

### Identification of OR open reading frames split across two exons

Having defined full-length gene models for a large portion of the mouse OR gene repertoire, we reanalysed the resulting transcripts to check if any showed evidence for full-length multi-exonic ORFs. We focused our analysis on the mouse repertoire since it contains a much higher number of full-length gene models. We used an ad hoc script (https://gitlab.com/olfr/olfr_multi_exon_orf_finder/tree/master) to identify all transcripts whose longest predicted ORF spanned multiple exons and encoded a protein greater than 300 aa. The resulting list of 202 different transcripts (171 genes) was filtered to remove transcripts that had a second conserved in-frame ATG downstream of the splice site that still generated a protein of more than 300 aa. We further removed any transcripts that had deletions in the seven transmembrane domains (assessed from a multiple alignment of all receptors) as these are likely to be pseudogenes. Finally, to increase the likelihood of retaining functional proteins, we only selected genes whose orthologues had a conserved starting methionine (i.e., in the same relative position). This procedure resulted in a list of 58 candidate transcripts with a split ORF. Each was manually curated to ensure that the predicted ORF satisfied all of the criteria described above to be biotyped as protein-coding. We further required good conservation of the splice junction in the orthologous loci in other mammals, or in the closest paralogues if an orthologue could not be found. The transcripts that satisfied all these requirements are detailed in Supplementary File 5, which also contains the corresponding human orthologues with conserved split ORF structures.

### Single-cell RNAseq of mature mouse olfactory sensory neurons

We dissected whole olfactory mucosa from two male and two female three-day old heterozygous OMP-GFP mice (B6;129P2-Omptm3Mom/MomJ, The Jackson Laboratory, Stock # 006667) (44) that were backcrossed 10 times with C57BL/6 animals, designated as OMP-GFP (B6-N11). The dissected tissue from all animals was pooled, minced and then enzymatically digested in HBSS without Ca^2+^ and Mg^2+^, supplemented with 44 U/mL dispase (Invitrogen), 1000 U/mL collagenase type II (Invitrogen) and 10 mg/mL DNaseI (Roche), for 15-20 min at 37°C with agitation. Digested tissue was centrifuged at 0.4 × 1000 rcf for 5 min and washed twice in HBSS without Ca^2+^ and Mg^2+^. The dissociated cell suspension was passed through a series of filters: 100 μm (Falcon), 70 μm (Flacon) and 20 μm cell strainers (pluriSelect). The final dissociated cells were resuspended in 0.5% BSA in PBS without Ca^2+^ and Mg^2+^. Single cells were isolated using a Nikon Narishige microinjection setup under a fluorescence Nikon TE300 microscope. GFP fluorescence was verified on a flat screen using NIS Elements v4.5 software (Nikon) in order to ensure only GFP-positive cells were selected. Isolated cells were washed twice and then pipetted into a 0.2 mL tube, which was immediately frozen on dry ice.

Single-cell cDNA libraries were prepared using SMART-Seq v4 (Clontech) according to the manufacturer’s recommendations. Briefly, single cells were lysed and reverse transcription was performed with SMART-Seq v4 Oligonucleotide and SMARTScribe Reverse Transcriptase to generate nearly full-length cDNA libraries. Libraries were purified twice with AMPure XP system (Beckman Coulter) to minimize primer dimers. The size of the libraries was checked on high-sensitivity DNA chips (Agilent); cells with abundant short cDNA (average size < 1.3 kb) were discarded. Purified cDNAs were used to construct libraries for sequencing using the Nextera XT DNA Library Preparation Kit (Illumina) according to manufacturer’s recommendations. Briefly, cDNAs were fragmented to ~300 bp, followed by PCR amplification with index primers. Libraries were purified with AMPure XP system (Beckman Coulter), normalised and pooled with different i7 indexes. Pooled libraries were sequenced on the Illumina HiSeq X Ten platform, to produce 150 bp paired-end fragments (Novogene Co., Beijing). Raw data have been deposited to Array Express (https://www.ebi.ac.uk/arrayexpress/) under accession number E-MTAB-8285.

Sequencing reads were mapped to the mouse reference genome (GRCm38) using STAR (45) version 2.6.0c, with parameters --outFilterMismatchNmax 6 –outFilterMatchNminOverLread 5 --outFilterScoreMinOverLread 0.5 --outSAMtype BAM SortedByCoordinate -- outFilterType BySJout --outFilterMultimapNmax 20 --alignSJoverhangMin 8 -- alignSJDBoverhangMin 1 --alignIntronMin 20 --alignIntronMax 1000000 --alignMatesGapMax 1000000 --outSAMstrandField intronMotif --quantMode GeneCounts. We guided the mapping with Ensembl annotation version 93 (46) that includes the Ensembl-HAVANA OR gene annotation. We enabled STAR’s quantification mode to obtain the number of fragments mapped to each gene.

All 34 cells but one showed good performance on quality-control statistics (library size, percent of reads mapping to mitochondrial reads, and number of genes detected); the failed sample had a much lower number of genes detected compared to all other samples and was removed from the analysis. Raw counts were normalised with the algorithm implemented in the Bioconductor package scran (47). For each single cell, we visually inspected the alignments of all OR genes expressed at 30 or more normalised counts. When clear mapping artefacts were detected, the counts of the corresponding OR gene were set to 0. Details of this procedure along with the code used to analyse the data can be found in https://github.com/xibarrasoria/ORgeneAnnotation_HAVANA.

### Phylogenetic analysis of protein-coding ORs

To reconstruct the phylogenetic relationship between ORs we first aligned the protein sequences of all protein-coding genes. For the human repertoire we used MUSCLE (48), which can handle up to 500 sequences (https://www.ebi.ac.uk/Tools/msa/muscle/; (49)), and for mouse we used CLUSTAL Omega (50), which aligns up to 2000 sequences (https://www.ebi.ac.uk/Tools/msa/clustalo/; (49)). A phylogenetic tree was constructed from the resulting multiple alignments using the BIONJ algorithm (51) (a modified neighbour-joining procedure), in Phylogeny.fr (52) (http://www.phylogeny.fr/one_task.cgi?task_type=bionj). The resulting trees were visualised with the Interactive Tree of Life tool (53) (http://itol.embl.de).

## RESULTS

### The OR gene repertoires in the human and mouse reference genomes

Most OR genes have previously been annotated *in silico,* by homology searches based on a small number of experimentally derived OR sequences, and often include only the coding region of the gene. In order to comprehensively annotate the OR gene repertoires of the human and mouse genomes we developed an expert curation pipeline (Figure 1; Methods) to identify, annotate, and refine the gene models for all OR genes. We identified 873 human and 1483 mouse loci encoding OR genes and pseudogenes (Table 1; Supplementary File 1). As previously reported (11–13), a typical OR gene consists of a short 5’ untranslated region (UTR) composed of one to six alternatively spliced non-coding exons, followed by a long exon containing the open reading frame (ORF) plus a substantial 3’ UTR (Figure 2).

**Table 1:** OR loci in the human and mouse genomes. The number of gene biotypes (protein-coding or pseudogenised), proportion of protein-coding genes containing an uORF (upstream open reading frame) and/or uATG (upstream methionine codon), and the subtype (unprocessed, polymorphic or unitary) of the pseudogenes are shown (mouse unitary pseudogenes were not determined) Also, the number of OR loci with exons that overlap the exon(s) of an adjacent gene on the same strand. Overlapping loci represent genes that either share 5’ UTR exon(s) or are readthrough transcripts predicting a chimeric protein.

|  | Human | Mouse |
| --- | --- | --- |
| **Total** | **873** | **1483** |
| **Protein-coding** | **389 (44.55%)** | **1141 (76.9%)** |
| uORF | 212 (54.5%) | 986 (86.4%) |
| uATG | 53 (13.6%) | 245 (21.5%) |
| **Pseudogene** | **485 (55.55%)** | **342 (23.1%)** |
| Unprocessed | 448 (92.4%) | 286 (83.6%) |
| Polymorphic | 19 (3.9%) | 56 (16.4%) |
| Unitary | 18 (3.7%) | NA |
| Overlapping loci | 29 | 54 |
| Shared 5' UTR exons | 7 | 7 |
| Chimeric protein | 11 | 11 |

To identify the OR genes with protein-coding potential, we manually assessed each locus based on the presence of: 1) an intact intronless ORF encoding a protein between 300 and 350 aa; 2) a predicted seven-transmembrane domain structure, which is characteristic of OR genes; 3) extracellular amino-terminal and intracellular carboxy-terminal domains; and 4) good cross-species conservation. Loci that failed at any of these criteria were annotated as pseudogenes and the *pseudogenic CDS* was defined as the region of the transcript with homology to the CDS of a functional OR protein. Many genes contained one or more in-frame upstream ATGs (Table 1) and we identified the ATG most likely to be used for initiation via conservation rather than taking the available longest ORF. Based on this, we changed the ORF length for 44 human and 90 mouse protein-coding OR genes.

The mouse genome had a much higher proportion of protein-coding loci (76.9%) compared to human (44.6%). The average length of the CDS for protein-coding genes was comparable in both species: 315.4 and 313.9 aa in human and mouse respectively. The *pseudogenic CDS* of pseudogenes was much larger in human (289.77 aa) than in mouse (220.09 aa), suggesting that OR gene losses occurred earlier in the mouse (Figure 2).

We compared our set of annotated OR genes and pseudogenes to the gene models present in other databases: RefSeq for both species (16), HORDE for human (18) and MGI for mouse (17). There were several discrepancies between the existing databases and our results, but these were much less prevalent for the human repertoire (1.4% loci were amended, compared to 9% for mouse), most likely due to HORDE’s extensive analysis and community feedback program. Based on our in-depth manual analysis we amended the biotype annotation of 9 human and 46 mouse genes, along with the identification of polymorphic pseudogenes and inclusion of completely novel loci, mostly pseudogenes (Table 2 and Supplementary File 1).

**Table 2:** Number of human and mouse OR genes with amended biotype annotation, and number of loci added or removed from the reference genome annotation.

| Amendment | Human | Mouse |
| --- | --- | --- |
| Pseudogene to protein-coding | 7 | 42 |
| Protein-coding to pseudogene | 2 | 4 |
| Polymorphic pseudogenes | 4 | 56 |
| Polymorphic to unprocessed pseudogene | 2 | 0 |
| Novel OR genes (pseudogenes) added | 2 (6) | 1 (50) |
| OR pseudogenes removed | 2 | 3 |

Additionally, we identified 41 OR loci present in the MGI database that could not be uniquely aligned to the reference mouse genome (Supplementary File 4), probably due to haplotypic differences and copy number variation between inbred mouse strains (8). Similarly, several OR loci were absent or incorrectly mapped on the reference human genome. For example, a recent human segmental duplication (chr15:21534404-22126421; GRCh38) was found to contain a duplicated cluster of nine OR loci. However, for eight of the nine duplicates, only one copy was annotated. We therefore added the missing eight paralogues, two of which were protein-coding (Supplementary File 1).

Finally, previous work has shown that a large proportion of the human OR protein-coding repertoire contains segregating pseudogenes in the population (10, 54). Some of these were annotated as unprocessed pseudogenes in the reference genome, but contain variation that resurrects them into protein-coding genes (5). We confirmed 16 such cases, previously annotated in HORDE, and we identified an additional 3 OR pseudogenes *(OR10J3, OR2T7, OR4C45)* resurrected by a single nucleotide polymorphism. To extend this analysis to the mouse repertoire, we mined variation data from the Mouse Genomes Project (27) and identified 56 polymorphic pseudogenes (OR pseudogenes in the reference annotation that contain protein-coding alleles in other mouse strains; Supplementary File 1).

In summary, we have comprehensively annotated the human and mouse OR gene repertoires, correcting errors from automated pipelines and unifying the criteria used to define gene biotypes and the coding sequence. In our view, this effort represents the most accurate catalogue of human and mouse OR genes available to date.

### The non-coding structure of OR genes

To define the UTR structure of OR genes, we performed reference-guided assembly of RNAseq data from human and mouse whole olfactory mucosa samples. For mouse, we used twelve samples from previous studies (11, 34). For human, we combined data from two independent studies (12, 35), and sequenced six additional samples to increase the coverage and representation of the OR genes (Methods). We visually examined each of the generated gene models in both species and manually curated them to remove artefacts and errors (Methods). We also considered evidence from available mRNAs, ESTs and PacBio sequences (55) from GenBank. Combined, these experimental data enabled the annotation of transcript models for 74% of human and 94% of mouse protein-coding OR loci (Figure 2). In contrast, only 17% of human and 12% of mouse OR pseudogenes were transcribed. These transcribed pseudogenes predominantly corresponded to gene models with minimally disrupted ORFs, suggesting they have been recently pseudogenised and still retain the regulatory elements for transcription. For the remaining OR loci, the number of sequencing reads was insufficient to confidently construct a gene model. Importantly, we note that a fraction of the OR gene models is likely to be incomplete due to low coverage from the RNAseq data. Indeed, when we grouped the human protein-coding OR genes by length we observed that the majority of genes containing only the CDS (< 1.1 kb) were expressed at very low levels in all samples, while those with gene models of >3 kb were expressed at moderate to high levels (Figure 3). Overall, OR genes ≤3 kb in length had significantly lower expression than their longer counterparts (Wilcoxon rank sum test, one-tail, p-value < 2.2e-16), suggesting that their shorter gene models are the result of insufficient transcriptional data to achieve full-length annotation. This observation was extended to the mouse repertoire (Supplementary Figure 1), despite the higher quality and coverage of mouse data.

**Figure 3.**
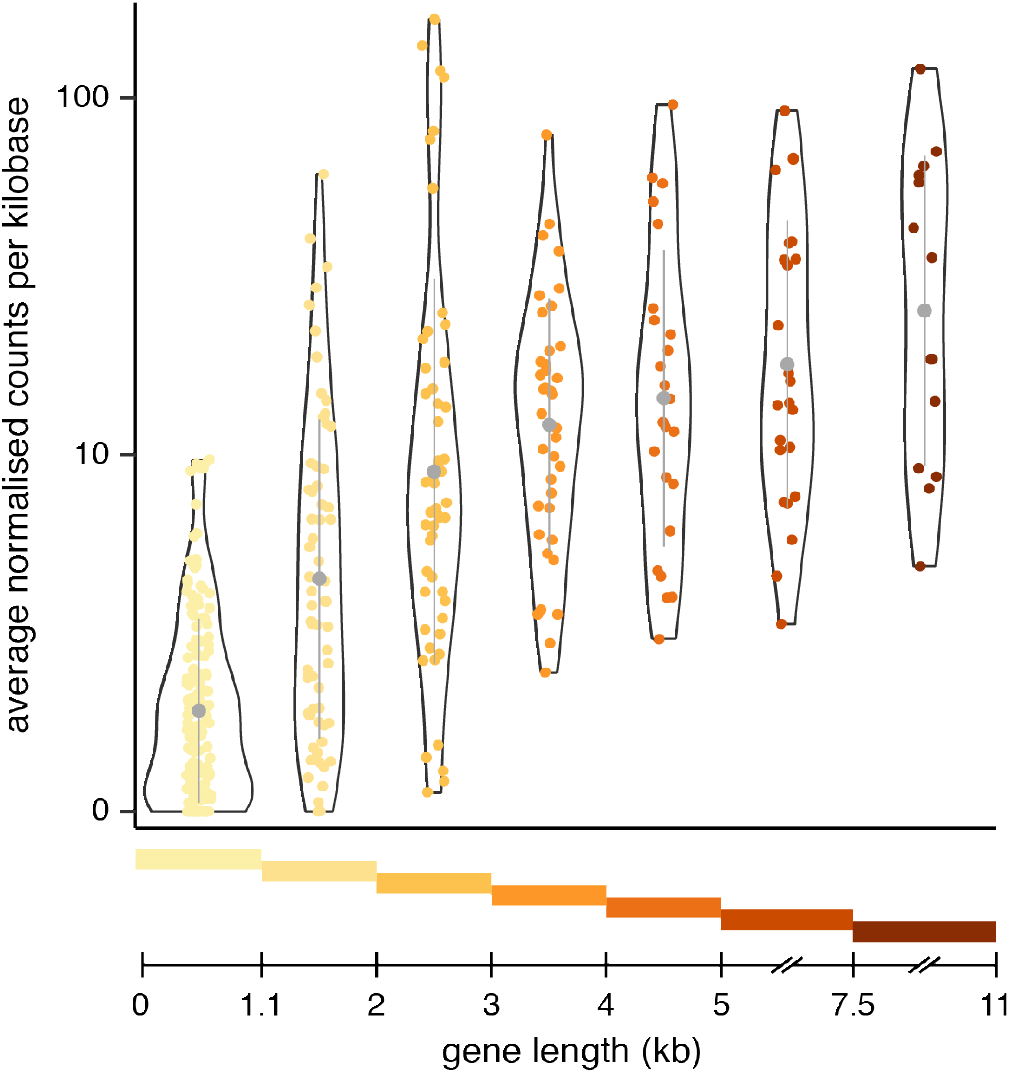
Short OR gene models are likely to be incomplete due to low expression levels. Violin plots of the mean expression levels for all human protein-coding OR genes grouped by length. The coloured bars at the bottom indicate the range of gene lengths included in each group. The median (circle) ± one standard deviation (vertical line) is indicated in grey. Expression levels are per kilobase (kb) of gene length. Genes with shorter gene models are expressed at significantly lower levels than those of 3 kb or larger, suggesting their models are incomplete.

The 5’ UTR was on average 192 bp for human and 391 bp for mouse OR protein-coding transcripts, and was formed by multiple short exons (Figure 2, Table 3). In both species, these 5’ UTR exons were frequently associated with alternative splicing, with most multi-exonic genes showing two or more alternative transcripts (60% for human and 55% for mouse). The majority (~62%) of OR genes had only two alternatively spliced transcripts, although some had up to nine different splice variants (Supplementary Figure 2). In contrast, the 3’ UTR was much larger, approximately 1.2 kb and 1.8 kb in human and mouse respectively (Figure 2, Table 3). A fraction of the OR loci in both species showed a drop in coverage across the distal region of the 3’ UTR, suggesting alternative polyadenylation sites. In these cases, we used the longest 3’ UTR supported by transcriptional data in our transcript models. However, a recent study (56) experimentally validated alternative polyadenylation sites for a fraction of the mouse OR gene repertoire, validating this phenomenon.

**Table 3:** Mean ± standard error for the longest transcript per locus.

|  | Human | Mouse |
| --- | --- | --- |
| **Protein-coding** |  |  |
| 5' UTR | 191.9 ± 17.32 bp | 390.74 ± 11.9 bp |
| CDS | 315.4 ± 0.43 aa | 313.9 ± 0.17 aa |
| 3' UTR | 1166.36 ± 91.53 bp | 1831.82 ± 46.68 bp |
| Number of exons | 1 to 6 (mean 1.69) | 1 to 7 (mean 2.33) |
| **Pseudogene** |  |  |
| 5' UTR | 46.1 ± 8.42 bp | 60.64 ± 26.32 bp |
| *Pseudogenic CDS* | 289.8 ± 3.02 aa | 220.1 ± 5.67 aa |
| 3' UTR | 154.27 ± 28.39 bp | 79.69 ± 21.24 bp |
| Number of exons | 1 to 5 (mean 1.21) | 1 to 5 (mean 1.36) |

We also identified a number of readthrough OR loci that shared the 5’ UTR exon(s) of an upstream gene which was frequently another OR gene (Table1, Figure 4, and Supplementary Figures 3-4). In all cases, the splice junction connecting the two genes was supported by transcriptional evidence from RNAseq and/or EST and mRNA sequences. Similarly, both human and mouse each contained 11 OR loci involved in chimeric transcripts (Table1). One chimeric transcript predicted an intact CDS and the remainder predicted either truncated ORFs or transcripts susceptible to degradation by nonsense-mediated decay. Finally, an additional 11 OR loci in human and 36 in mouse overlapped with at least one other gene on the same strand (Table 1). Most of these were remnants of OR pseudogenes completely embedded within the 3’ UTR of protein-coding genes.

**Figure 4.**
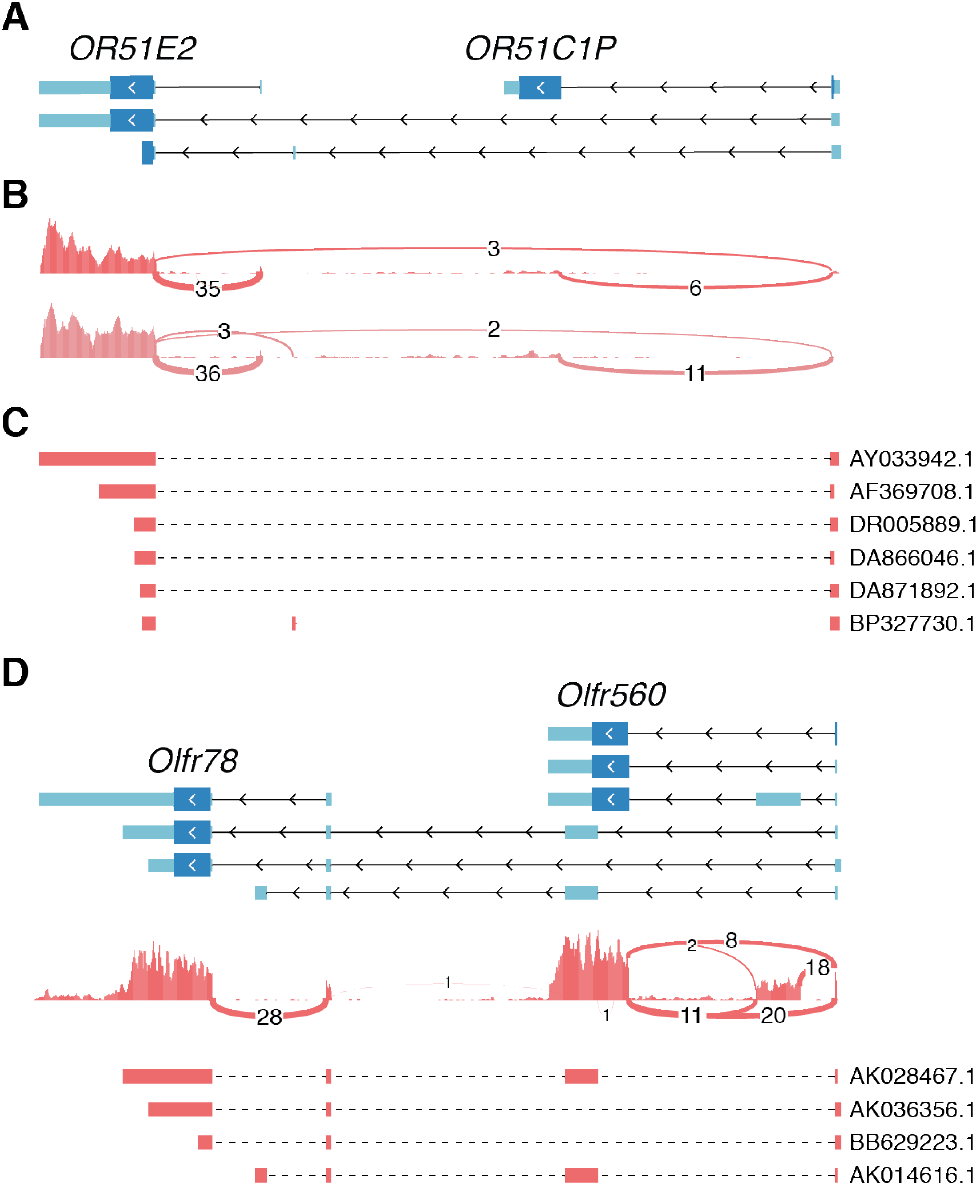
Some OR genes share 5’ UTR exons. **(A)** Transcript models for two human OR genes. Exons are depicted as boxes and introns as connecting lines. The arrowheads indicate the direction of transcription. The coding sequence is represented by taller, darker blue boxes. OR51E2 contains transcripts that splice across to the most 5’ exon of the adjacent OR51C1P gene. **(B)** Coverage plot of the aggregated RNA-seq reads from this study (top; 9 samples) and from Olender et al. (bottom; 4 samples). Lines represent splice junctions and the number of supporting reads are indicated. **(C)** Further support for the splice junction spanning both genes can be found in several mRNA and EST clones deposited to GenBank; accession numbers are indicated. For each sequence, the red boxes represent alignments to the reference genome. **(D)** Same as in A-C but for the orthologous genes in the mouse genome. The sharing of 5’ UTR exons is supported by mRNA and EST clones from GenBank (derived from non-olfactory tissues), but is not observed in the RNAseq data from olfactory mucosa. The coverage plot is from the mouse RNA-seq data of all 12 samples together.

As noted previously (13), a large proportion of the protein-coding OR genes had additional ORFs upstream of the iATG (referred to as uORFs): 54.6% in human and 86.1% in mouse (Table 1). A lower fraction had an in-frame uATG, 13.6% and 21.5% of the human and mouse protein-coding repertoires, respectively (Table 1). Both uORFs and uATGs have been shown to downregulate translation (57, 58).

### Protein-coding OR genes with coding sequences split across two exons

We have previously reported some mouse OR transcripts contain a predicted intact ORF encoded across two exons (11). We therefore analysed all mouse OR transcripts to identify potential full-length ORFs interrupted by an intron (Methods). Only cases where the initiation methionine and splice junction were conserved in the orthologous sequences of other mammals were considered; for OR genes that lacked orthologues the closest paralogues were used instead.

We identified 47 mouse OR transcripts (from 41 genes) satisfying these criteria (Supplementary File 5), which we will refer to as *split* OR genes (Figure 5 and Supplementary Figure 5). Nine of these mouse split OR genes had an orthologous split OR structure in human. We identified an additional four split OR genes in the human repertoire that lacked a mouse orthologue, bringing the total of human split OR genes identified to 13 (Supplementary File 5).

**Figure 5.**
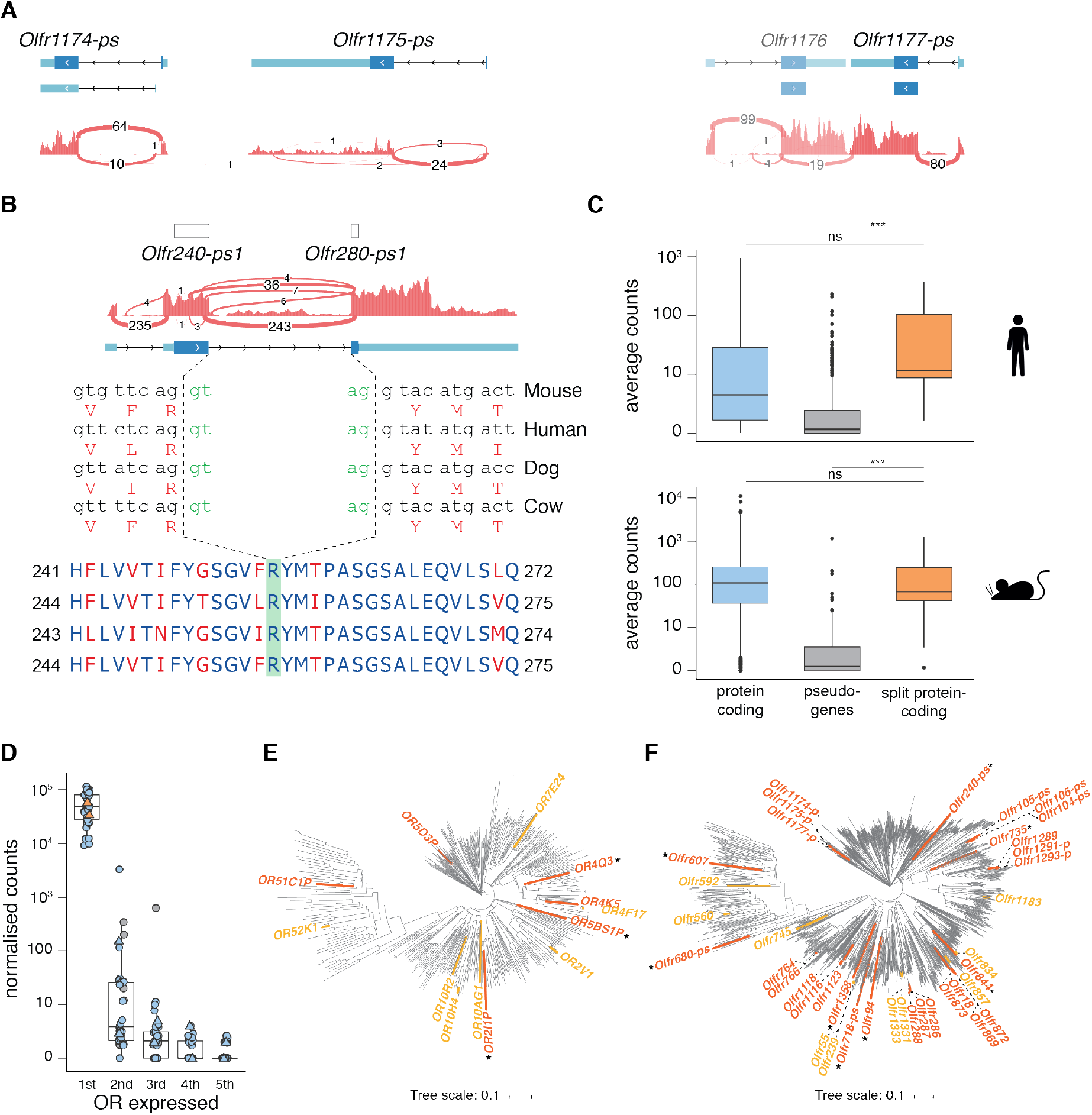
Some OR genes have an intron within the coding sequence. **(A)** Example of three mouse OR loci (*Olfr1174-ps, Olfr1175-ps and Olfr1177-ps*) previously annotated as pseudogenes due to the lack of a conserved Initiation ATG. However, an IATG can be found In the upstream exon, producing an Intact open reading frame of ~320 amino acids. Below the gene models, the coverage track for the combined RNAseq data from all mouse samples (n=12) Is shown, along with the number of reads supporting each exon junction. **(B)** At the top, previous annotation showed truncated open reading frames (ORFs) of two OR pseudogenes. We Identify an Intact ORF spanning the two loci with strong support from the RNAseq data. Below Is the nucleotide sequence around the splice junction, for mouse and the corresponding orthologues In other mammals; the splice junction Is conserved. At the bottom Is the multiple alignment of the protein sequences for the same species; conserved amino acids are In blue, variable ones In red. **(C)** Boxplots of the average normalised counts for all protein-coding and pseudogene OR genes, plus those that become protein coding after considering ORFs containing an Intron (split), for human (top) and mouse (bottom). The split OR genes are expressed at similar levels to protein-coding loci, and significantly higher than pseudogenes. Wllcoxon rank sum test, one-tall, ns = not significant; *** p-value < 1×10-7.**(D)** Boxplots of the normalised expression levels of the five most highly expressed OR genes In each of 33 single mature OSNs. The values for each cell are Indicated, coloured following the same scheme as In C. Values for the cells expressing split OR genes are shown as triangles. All 33 mature OSNs express one OR gene at very high levels (1st); expression then drops rapidly by more than three orders of magnitude, supporting monogenic OR expression. The split OR genes (orange) are expressed at the same levels as the other protein-coding genes, and can similarly Induce monogenic expression. **(E)** Phylogenetic tree of all protein-coding human OR genes. The split OR genes are highlighted In orange and yellow; the latter correspond to genes with two protein-coding Isoforms, one split ORF and the other ORF contained within a single exon. Asterisks Indicate ORs within clades that previously contained only pseudogenes. **(F)** Same as E but for the mouse protein-coding OR genes.

In both species, the split OR genes were scattered across the genome, found in 7 and 10 different chromosomes in human and mouse, respectively. In >90% of the split OR genes (44/47 in mouse and 12/13 in human), the intron was inserted into the extracellular N-terminal domain or within the TM1 region. The average size of this intron was 3841.3 bp (range 1384-7585 bp) for human and 3413.1 bp (range 550-22628 bp) for mouse (Supplementary File 5), and this was not significantly different from the length of the most 3’ intron of OR genes with their CDS contained within a single exon (the most 3’ intron is generally the intron preceding the CDS; Wilcoxon rank sum test, two-tails, p-value = 0.2847). We could not identify any distinct sequence features in the intron sequences of the split OR genes compared to their intronless counterparts, including repeat element composition.

We observed two classes of split OR transcripts. The first consists of loci previously biotyped as pseudogenes because they lacked a conserved iATG or N-terminal domain (Figure 5A). These features were recovered in the adjacent exon and were subsequently amended to protein-coding. The second class comprises loci with alternatively spliced transcripts, some with the ORF contained within a single exon whilst others have the ORF split across two exons. These represent OR genes encoding isoforms with variable N-terminal domains. Interestingly, we also found a locus in the mouse genome with two annotated OR pseudogenes that, upon inspection of the RNAseq data, revealed a single gene (ENSMUST00000216180.1) with an intact coding sequence split across two exons, interrupted by repetitive sequences (Figure 5B). The human orthologue *(OR5BS1P),* as well as orthologues from other mammals, all showed the same intact ORF split across exons.

Split OR genes were expressed at similar levels to protein-coding genes but significantly higher than pseudogenes (Wilcoxon rank sum test, one-tail, p-value < 1×10-7; Figure 5C) suggesting that the split OR genes may encode functional OR proteins. We reasoned that one way to assess whether split OR genes are functional would be to identify single mature OSNs that express a single split OR gene at high levels (59, 60). To this end, we performed singlecell RNAseq on 33 manually picked single GFP-expressing OSNs from heterozygous OMP-GFP gene-targeted mice (44), with OMP being a marker for mature OSNs. Each of these 33 OSNs expressed a different OR gene abundantly and, generally, this OR gene was within the top five most highly expressed genes in the cell (~53,910 ± 31,099.2 normalised counts; mean ± standard deviation; Figure 5D). Interestingly, two of the 33 OSNs expressed a split OR gene *(Olfr718-ps1* or *Olfr766)* at levels comparable to those of intronless OR genes in the other 31 OSNs, and the levels of the second highest expressed OR genes were hundreds to thousands times lower (Figure 5D). Thus, the two cells expressing *Olfr718-ps1* and *Olfr766* were indistinguishable from the other 31 OSNs in terms of expression in the OR gene repertoire, suggesting that they encode functional OR proteins.

Finally, in order to determine the evolutionary history of the split OR genes, we built phylogenetic trees for all human and mouse protein-coding OR sequences (Figure 5E-F). In mouse, there were 23 independent insertion events of an intron into the ORF. Nine of these occurred within phylogenetically related small clusters of OR genes (two to four members), all with a common single-exon OR ancestor gene that gained the intron. Importantly, eight of the split OR genes belonged to clades initially believed to encompass only pseudogenes. All the human split OR loci were phylogenetically independent events.

## DISCUSSION

In this study, we have assessed every OR locus in the human and mouse reference genomes, taking great care to ensure consistency of their annotation within and between species. For each OR gene we have considered structural features as well as protein conservation across mammals to determine protein-coding potential and to define the coding sequence. The Ensembl-HAVANA curation project therefore provides the most comprehensive catalogue of OR genes in human and mouse, with a uniform set of criteria determining biotype and feature annotation of all OR genes.

### Annotation of full-length OR transcript models

Accurate and complete annotation of the OR gene structure requires transcriptomic data. In the last few years several datasets have been generated from mouse olfactory mucosa samples and used to reconstruct full-length OR transcripts (11, 13). However, a lack of integration of these results into genome browsers and their associated databases has limited their dissemination and inclusion into olfactory studies. We have now evaluated the transcript models generated by in silico assembly of the RNAseq data to generate a high-quality set of expertly curated gene models. We have included these into the GENCODE reference gene set, making them widely available to the research community.

Annotation of the human OR gene repertoire has proved more challenging, mostly due to the difficulty in obtaining high-quality olfactory mucosa samples from humans. To date, there are very few studies that have sequenced the transcriptome of human olfactory mucosa samples (12, 35). Olender at al. (12) annotated transcript models for a hundred OR genes, representing only 11.5% of the complete repertoire from human biopsy samples. We have produced six additional high-quality transcriptomes from human surgical samples of olfactory mucosa, and combined with the previous resources, generated extended gene models for 254 OR genes, including 49.9% of the protein-coding OR loci, a ~2.5-fold increase. Further evidence from mRNA and EST clones supported gene models for an additional ~25% of the protein-coding OR loci, which lacked RNAseq data. Nonetheless, many of the models are still incomplete and contain limited representation of the untranslated regions, with only around a quarter of the protein-coding models longer than 3kb. The insufficient data for most OR genes is due to their low representation in whole-tissue transcriptomes. In order to obtain full-length gene models, it will be necessary not only to increase the number of human samples available, but to implement targeted approaches (such as (55, 61)) that can enrich for OR transcripts specifically.

The fraction of the repertoire with full-length gene models suggests some interesting observations. First, there are several cases of 5’ UTR exon sharing with neighbouring genes (Figure 4 and Supplementary Figures 3-4). Some of these, however, are only supported by mRNA and EST data that was obtained from non-olfactory tissues, and there is no evidence of exon sharing in the RNAseq data from olfactory mucosa samples (Figure 4D). This observation suggests that some of the transcripts involving two OR genes might be expressed in a tissue-specific manner, outside the olfactory system. For example, *OR51E2,* which contains an isoform including the 5’ UTR exon from *OR51C1P* (Figure 4A), is highly expressed in prostate epithelium and has been identified as a biomarker for prostate cancer (62). Secondly, we observed alternatively spliced transcripts with an intron truncating the ORF. And, lastly, we also noticed alternatively spliced transcripts that completely skip the coding exon, resulting in transcripts that contain only non-coding sequence. All these might be involved in regulation of OR expression and/or translation.

### OR genes encoded across two exons

Contrary to the dogma held in the field since 1991 that vertebrate OR genes are encoded within a single exon (1), we have identified 13 human and 41 mouse OR genes with a coding region split across two exons. Some of these loci only satisfy all criteria to be considered protein-coding when the two exons are taken into account and have thus been considered pseudogenes until now. Most of the cases we identified are well conserved across mammalian evolution; such strong evidence of purifying selection together with their high, protein-coding-like expression levels, support their classification as protein-coding. Furthermore, we found two of 33 single mature OSNs expressed a split OR gene at high levels, similar to levels of the OR genes with intronless coding regions expressed in the other 31 single mature OSNs. Taken together, these data strongly support that split OR genes encode functional receptor proteins, equivalent to intronless OR genes.

It is very likely that there are many other split OR genes in both the human and mouse genomes that remain unannotated due to our stringent filtering criteria. We have excluded any cases where a downstream highly conserved methionine produces an ORF of correct length coded within a single exon; it is still possible that the split ORF is translated, but proteomic data will be necessary to confirm this. Additionally, in many cases the sequence encoded in the upstream exon is very short, which makes it difficult to identify the orthologous region in other mammals to establish whether the starting methionine and splice junction are conserved. In these cases, we rejected the split isoform. Finally, we identified a set of pseudogenes only lacking part of the N-terminal domain; these could be split OR loci but since they lack transcriptional data we have not been able to define upstream exons.

Also intriguing is that several of the genes with a split ORF have an alternative isoform encoded within a single exon. Thus, these split OR genes contain two different protein isoforms, with slightly different N-terminal domains. Although some of these differ by only a few amino acids (three to 15), several have stretches of around 30 amino acids that are not common between the two isoforms. One of these genes, *Olfr55,* was captured in two mature OSNs previously sequenced through single-cell transcriptomic approaches (64). In these mature OSNs both transcript isoforms are monoallelically expressed at high levels within a single neuron, raising the possibility that both proteins are translated. Further work will be necessary to establish the functional relevance of this phenomenon.

## ACKNOWLEDGEMENTS

We would like to thank the Ensembl-HAVANA team for their contribution towards some of the initial OR annotation, Andrew E. Berry and Mark Thomas for help with the OR analysis.

## Author Contributions

A.F. conceived the project. L.V.G., M.J., M.K., and P.M. provided human samples. I.H.A.B. curated every OR locus and associated transcript models. S.F. generated Cufflinks transcript models. I.H.A.B., S.F., C.D., M.P.H., and D.P. contributed to validating and amending Cufflinks models. S.F. and J.M.G performed bioinformatic analyses. Z.Z. and M.K. generated single-cell RNAseq data. X.I-S. generated and analysed bulk and single-cell RNAseq data. I.H.A.B. and X.I-S. analysed data and generated figures. I.H.A.B. and X.I-S. wrote the manuscript. I.H.A.B., X.I-S., and A.F. edited the manuscript. J.H. and D.W.L. acquired funding for this study. A.F. and D.W.L. supervised the study. All authors read and approved the final manuscript.

## SUPPLEMENTARY MATERIALS

**Supplementary Figure 1 |** Same as Figure 3 but for mouse protein-coding genes.

**Supplementary Figure 2 | ORs have several isoforms per gene.** Barplots of the number of genes with the indicated number of different transcript isoforms. Genes have been split into protein-coding (top) and pseudogenes (bottom), and by species (human on the right, mouse on the left).

**Supplementary Figure 3 |**Same as Figure 4 but for the additional mouse OR genes that share a 5’ UTR exon with a neighbouring gene. mRNA, EST or PacBio clones supporting splice junctions between the two genes are indicated above the corresponding transcript.

**Supplementary Figure 4 |** Same as Figure 4 but for the additional human OR genes that share a 5’ UTR exon with a neighbouring gene. mRNA, EST or PacBio clones supporting splice junctions between the two genes are indicated above the corresponding transcript.

**Supplementary Figure 5 |** Additional example of a split OR gene. On chromosome 7, *Olfr682-ps1* was annotated as a pseudogene, but we identified an open reading frame (ORF) spanning two exons that codes for a 311 aa protein. This gene is a polymorphic pseudogene that, in the reference genome, contains a frameshift in the C-terminal domain (purple transcript); however, several mouse strains contain a 2bp indel at position 105,126,541 that restores the correct frame. The splice junction and protein sequence are conserved in several mammals, including dog, cow and sheep. *Olfr682-ps1* has a close paralogue, *Olfr680-ps1*, which shares 97% identity at the protein level. Whereas *Olfr680-ps1* lacks transcriptional evidence, we used the conservation with *Olfr682-ps1* and other mammals to annotate a full-length split transcript structure.

**Supplementary File 1 |** Table with all the human and mouse OR genes in the Ensembl-HAVANA annotation. For each gene we provide their official gene symbol, chromosomal location, description, biotype and associated Ensembl identifier (version 98).

**Supplementary File 2 |** GTF annotation file with all curated human OR transcript models from the Ensembl-HAVANA annotation.

**Supplementary File 3 |** GTF annotation file with all curated mouse OR transcript models from the Ensembl-HAVANA annotation.

**Supplementary File 4 |** List of mouse OR genes present in MGI that cannot be placed in the mouse reference genome.

**Supplementary File 5 |** List of curated human and mouse split OR genes.

**Supplementary File 6 |** Metadata for the human samples analysed, and normalised counts for human OR genes.

## REFERENCES

1. Buck L, Axel R. A novel multigene family may encode odorant receptors: a molecular basis for odor recognition. Cell. 1991;65(1):175–87.

2. Malnic B, Hirono J, Sato T, Buck LB. Combinatorial receptor codes for odors. Cell. 1999;96(5):713–23.

3. Glusman G, Yanai I, Rubin I, Lancet D. The complete human olfactory subgenome. Genome Res. 2001;11(5):685–702.

4. Zhang X, Firestein S. The olfactory receptor gene superfamily of the mouse. Nat Neurosci. 2002;5(2):124–33.

5. Olender T, Nativ N, Lancet D. HORDE: comprehensive resource for olfactory receptor genomics. Methods Mol Biol. 2013;1003:23–38.

6. Niimura Y, Matsui A, Touhara K. Extreme expansion of the olfactory receptor gene repertoire in African elephants and evolutionary dynamics of orthologous gene groups in 13 placental mammals. Genome Res. 2014;24(9):1485–96.

7. Young JM, Friedman C, Williams EM, Ross JA, Tonnes-Priddy L, Trask BJ. Different evolutionary processes shaped the mouse and human olfactory receptor gene families. Hum Mol Genet. 2002;11(5):535–46.

8. Lilue J, Doran AG, Fiddes IT, Abrudan M, Armstrong J, Bennett R, et al. Sixteen diverse laboratory mouse reference genomes define strain-specific haplotypes and novel functional loci. Nat Genet. 2018;50(11):1574–83.

9. Mainland JD, Keller A, Li YR, Zhou T, Trimmer C, Snyder LL, et al. The missense of smell: functional variability in the human odorant receptor repertoire. Nat Neurosci. 2014;17(1):114–20.

10. Olender T, Waszak SM, Viavant M, Khen M, Ben-Asher E, Reyes A, et al. Personal receptor repertoires: olfaction as a model. BMC Genomics. 2012;13:414.

11. Ibarra-Soria X, Levitin MO, Saraiva LR, Logan DW. The olfactory transcriptomes of mice. PLoS Genet. 2014;10(9):e1004593.

12. Olender T, Keydar I, Pinto JM, Tatarskyy P, Alkelai A, Chien MS, et al. The human olfactory transcriptome. BMC Genomics. 2016;17(1):619.

13. Shum EY, Espinoza JL, Ramaiah M, Wilkinson MF. Identification of novel post-transcriptional features in olfactory receptor family mRNAs. Nucleic Acids Res. 2015;43(19):9314–26.

14. Loveland JE, Gilbert JG, Griffiths E, Harrow JL. Community gene annotation in practice. Database (Oxford). 2012;2012:bas009.

15. Searle SM, Gilbert J, Iyer V, Clamp M. The otter annotation system. Genome Res. 2004;14(5):963–70.

16. O’Leary NA, Wright MW, Brister JR, Ciufo S, Haddad D, McVeigh R, et al. Reference sequence (RefSeq) database at NCBI: current status, taxonomic expansion, and functional annotation. Nucleic Acids Res. 2016;44(D1):D733–45.

17. Smith CL, Blake JA, Kadin JA, Richardson JE, Bult CJ, Mouse Genome Database G. Mouse Genome Database (MGD)-2018: knowledgebase for the laboratory mouse. Nucleic Acids Res. 2018;46(D1):D836–D42.

18. Safran M, Chalifa-Caspi V, Shmueli O, Olender T, Lapidot M, Rosen N, et al. Human Gene-Centric Databases at the Weizmann Institute of Science: GeneCards, UDB, CroW 21 and HORDE. Nucleic Acids Res. 2003;31(1):142–6.

19. Olender T, Feldmesser E, Atarot T, Eisenstein M, Lancet D. The olfactory receptor universe--from whole genome analysis to structure and evolution. Genet Mol Res. 2004;3(4):545–53.

20. Sonnhammer EL, Durbin R. A dot-matrix program with dynamic threshold control suited for genomic DNA and protein sequence analysis. Gene. 1995;167(1-2):GC1–10.

21. Kent WJ. BLAT--the BLAST-like alignment tool. Genome Res. 2002;12(4):656–64.

22. Tyner C, Barber GP, Casper J, Clawson H, Diekhans M, Eisenhart C, et al. The UCSC Genome Browser database: 2017 update. Nucleic Acids Res. 2017;45(D1):D626–D34.

23. Tusnady GE, Simon I. The HMMTOP transmembrane topology prediction server. Bioinformatics. 2001;17(9):849–50.

24. Hofmann K, Stoffel W. TMbase - A database of membrane spanning proteins segments. Biol Chem Hoppe-Seyler. 1993;374(166).

25. Krogh A, Larsson B, von Heijne G, Sonnhammer EL. Predicting transmembrane protein topology with a hidden Markov model: application to complete genomes. J Mol Biol. 2001;305(3):567–80.

26. Sherry ST, Ward MH, Kholodov M, Baker J, Phan L, Smigielski EM, et al. dbSNP: the NCBI database of genetic variation. Nucleic Acids Res. 2001;29(1):308–11.

27. Adams DJ, Doran AG, Lilue J, Keane TM. The Mouse Genomes Project: a repository of inbred laboratory mouse strain genomes. Mamm Genome. 2015;26(9-10):403–12.

28. Barson G, Griffiths E. SeqTools: visual tools for manual analysis of sequence alignments. BMC Res Notes. 2016;9:39.

29. Robinson JT, Thorvaldsdottir H, Winckler W, Guttman M, Lander ES, Getz G, et al. Integrative genomics viewer. Nature biotechnology. 2011;29(1):24–6.

30. Thorvaldsdottir H, Robinson JT, Mesirov JP. Integrative Genomics Viewer (IGV): high-performance genomics data visualization and exploration. Brief Bioinform. 2013;14(2):178–92.

31. Kokocinski F, Harrow J, Hubbard T. AnnoTrack--a tracking system for genome annotation. BMC Genomics. 2010;11:538.

32. Frankish A, Diekhans M, Ferreira AM, Johnson R, Jungreis I, Loveland J, et al. GENCODE reference annotation for the human and mouse genomes. Nucleic Acids Res. 2019;47(D1):D766–D73.

33. Harrow J, Frankish A, Gonzalez JM, Tapanari E, Diekhans M, Kokocinski F, et al. GENCODE: the reference human genome annotation for The ENCODE Project. Genome Res. 2012;22(9):1760–74.

34. Ibarra-Soria X, Nakahara TS, Lilue J, Jiang Y, Trimmer C, Souza MA, et al. Variation in olfactory neuron repertoires is genetically controlled and environmentally modulated. Elife. 2017;6:e21476.

35. Saraiva LR, Riveros-McKay F, Mezzavilla M, Abou-Moussa EH, Arayata CJ, Makhlouf M, et al. A transcriptomic atlas of mammalian olfactory mucosae reveals an evolutionary influence on food odor detection in humans. Sci Adv. 2019;5(7):eaax0396.

36. Kim D, Pertea G, Trapnell C, Pimentel H, Kelley R, Salzberg SL. TopHat2: accurate alignment of transcriptomes in the presence of insertions, deletions and gene fusions. Genome Biol. 2013;14(4):R36.

37. Trapnell C, Williams BA, Pertea G, Mortazavi A, Kwan G, van Baren MJ, et al. Transcript assembly and quantification by RNAseq reveals unannotated transcripts and isoform switching during cell differentiation. Nature biotechnology. 2010;28(5):511–5.

38. Yates A, Akanni W, Amode MR, Barrell D, Billis K, Carvalho-Silva D, et al. Ensembl 2016. Nucleic Acids Res. 2016;44(D1):D710–6.

39. Anders S, Pyl PT, Huber W. HTSeq-- a Python framework to work with high-throughput sequencing data. Bioinformatics. 2015;31(2):166–9.

40. Cunningham F, Achuthan P, Akanni W, Allen J, Amode MR, Armean IM, et al. Ensembl 2019. Nucleic Acids Res. 2019;47(D1]):D745–D51.

41. Love MI, Huber W, Anders S. Moderated estimation of fold change and dispersion for RNAseq data with DESeq2. Genome Biol. 2014;15(12):550.

42. Khan M, Vaes E, Mombaerts P. Temporal patterns of odorant receptor gene expression in adult and aged mice. Molecular and cellular neurosciences. 2013;57:120–9.

43. Team RC. R: A Language and Environment for Statistical Computing. 2014.

44. Potter SM, Zheng C, Koos DS, Feinstein P, Fraser SE, Mombaerts P. Structure and emergence of specific olfactory glomeruli in the mouse. J Neurosci. 2001;21(24):9713–23.

45. Dobin A, Davis CA, Schlesinger F, Drenkow J, Zaleski C, Jha S, et al. STAR: ultrafast universal RNAseq aligner. Bioinformatics. 2012;29(1):15–21.

46. Zerbino DR, Achuthan P, Akanni W, Amode MR, Barrell D, Bhai J, et al. Ensembl 2018. Nucleic Acids Res. 2018;46(D1):D754–D61.

47. Lun AT, McCarthy DJ, Marioni JC. A step-by-step workflow for low-level analysis of single-cell RNAseq data with Bioconductor. F1000Res. 2016;5:2122.

48. Edgar RC. MUSCLE: multiple sequence alignment with high accuracy and high throughput. Nucleic Acids Res. 2004;32(5):1792–7.

49. Chojnacki S, Cowley A, Lee J, Foix A, Lopez R. Programmatic access to bioinformatics tools from EMBL-EBI update: 2017. Nucleic Acids Res. 2017;45(W1):W550–W3.

50. Sievers F, Wilm A, Dineen D, Gibson TJ, Karplus K, Li W, et al. Fast, scalable generation of high-quality protein multiple sequence alignments using Clustal Omega. Mol Syst Biol. 2011;7:539.

51. Gascuel O. BIONJ: an improved version of the NJ algorithm based on a simple model of sequence data. Mol Biol Evol. 1997;14(7):685–95.

52. Dereeper A, Guignon V, Blanc G, Audic S, Buffet S, Chevenet F, et al. Phylogeny.fr: robust phylogenetic analysis for the non-specialist. Nucleic Acids Res. 2008;36(Web Server issue):W465–9.

53. Letunic I, Bork P. Interactive Tree Of Life (iTOL) v4: recent updates and new developments. Nucleic Acids Res. 2019;47(W1):W256–W9.

54. Menashe I, Man O, Lancet D, Gilad Y. Different noses for different people. Nat Genet. 2003;34(2):143–4.

55. Lagarde J, Uszczynska-Ratajczak B, Carbonell S, Perez-Lluch S, Abad A, Davis C, et al. High-throughput annotation of full-length long noncoding RNAs with capture long-read sequencing. Nat Genet. 2017;49(12):1731–40.

56. Doulazmi M, Cros C, Dusart I, Trembleau A, Dubacq C. Alternative polyadenylation produces multiple 3’ untranslated regions of odorant receptor mRNAs in mouse olfactory sensory neurons. BMC Genomics. 2019;20(1):577.

57. Kumar M, Srinivas V, Patankar S. Upstream AUGs and upstream ORFs can regulate the downstream ORF in Plasmodium falciparum. Malar J. 2015;14:512.

58. Zhang H, Wang Y, Lu J. Function and Evolution of Upstream ORFs in Eukaryotes. Trends Biochem Sci. 2019.

59. Lewcock JW, Reed RR. A feedback mechanism regulates monoallelic odorant receptor expression. Proc Natl Acad Sci U S A. 2004;101(4):1069–74.

60. Shykind BM, Rohani SC, O’Donnell S, Nemes A, Mendelsohn M, Sun Y, et al. Gene switching and the stability of odorant receptor gene choice. Cell. 2004;117(6):801–15.

61. Sheynkman GM, Tuttle KS, Tseng E, Underwood JG, Yu L, Dong D, et al. ORF Capture-Seq: a versatile method for targeted identification of full-length isoforms. bioRxiv. 2019:604157.

62. Rodriguez M, Siwko S, Liu M. Prostate-Specific G-Protein Coupled Receptor, an Emerging Biomarker Regulating Inflammation and Prostate Cancer Invasion. Curr Mol Med. 2016;16(6):526–32.

63. Dalton RP, Lyons DB, Lomvardas S. Co-opting the unfolded protein response to elicit olfactory receptor feedback. Cell. 2013;155(2):321–32.

64. Saraiva LR, Ibarra-Soria X, Khan M, Omura M, Scialdone A, Mombaerts P, et al. Hierarchical deconstruction of mouse olfactory sensory neurons: from whole mucosa to single-cell RNAseq. Sci Rep. 2015;5:18178.

