## Supplementary Figures for "Expert Curation of the Human and Mouse Olfactory Receptor Gene Repertoires Identifies Conserved Coding Regions Split Across Two Exons"

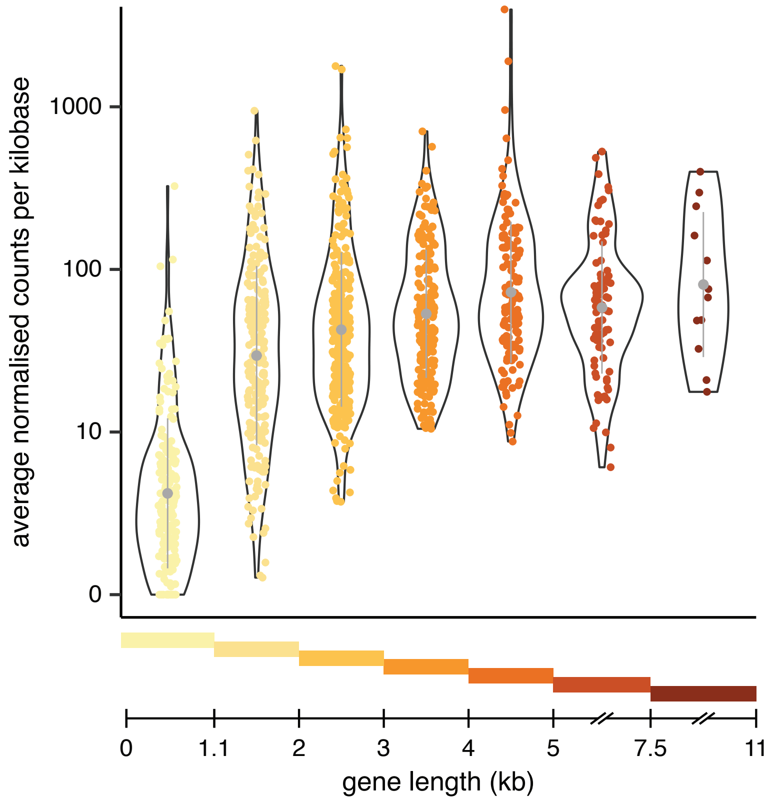


**Supplementary Figure 1 I** Same as Figure 3 but for mouse protein-coding genes.

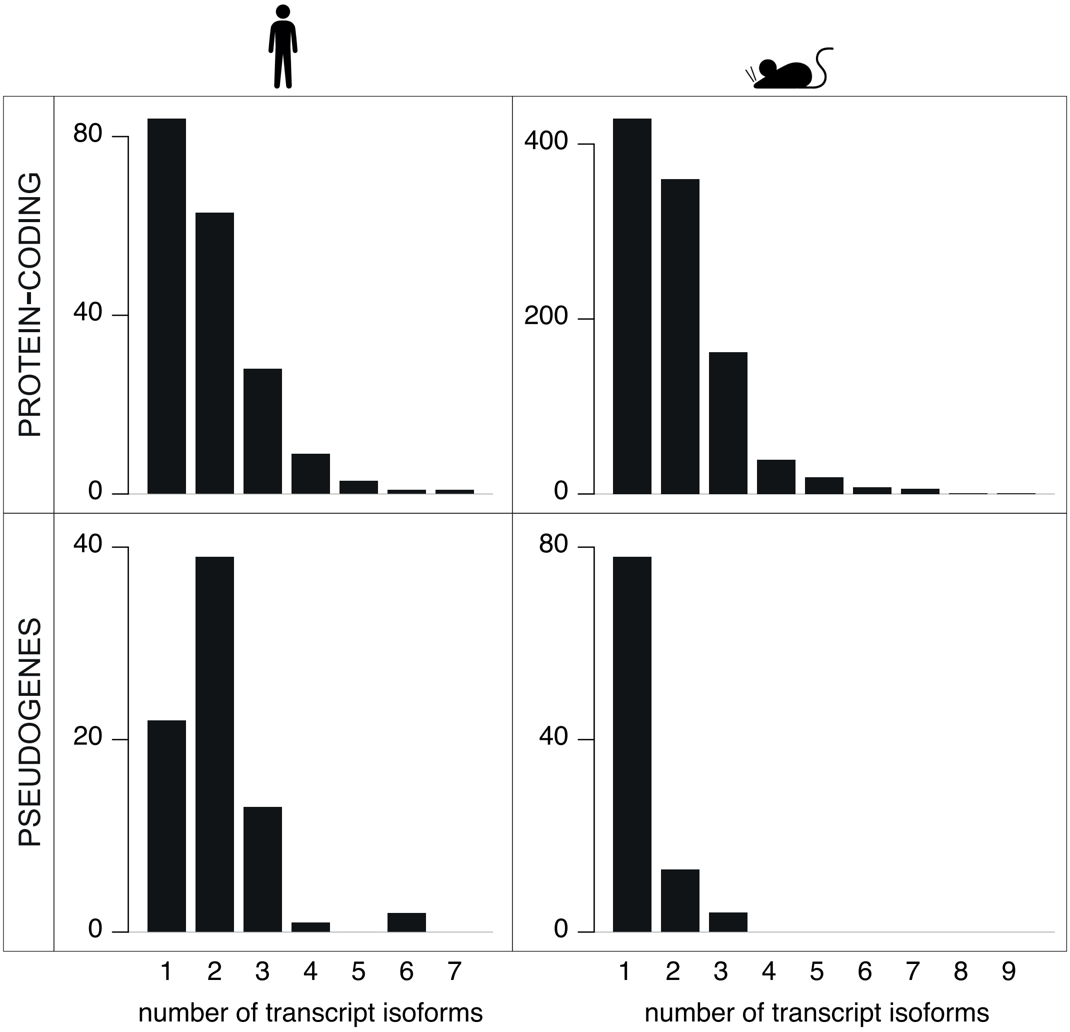


**Supplementary Figure 2 I Olfactory receptors have several isoforms per gene.** Barplots of the number of genes with the indicated number of different transcript isoforms. Genes have been split into protein-coding (top) and pseudogenes (bottom), and by species (human on the right, mouse on the left).

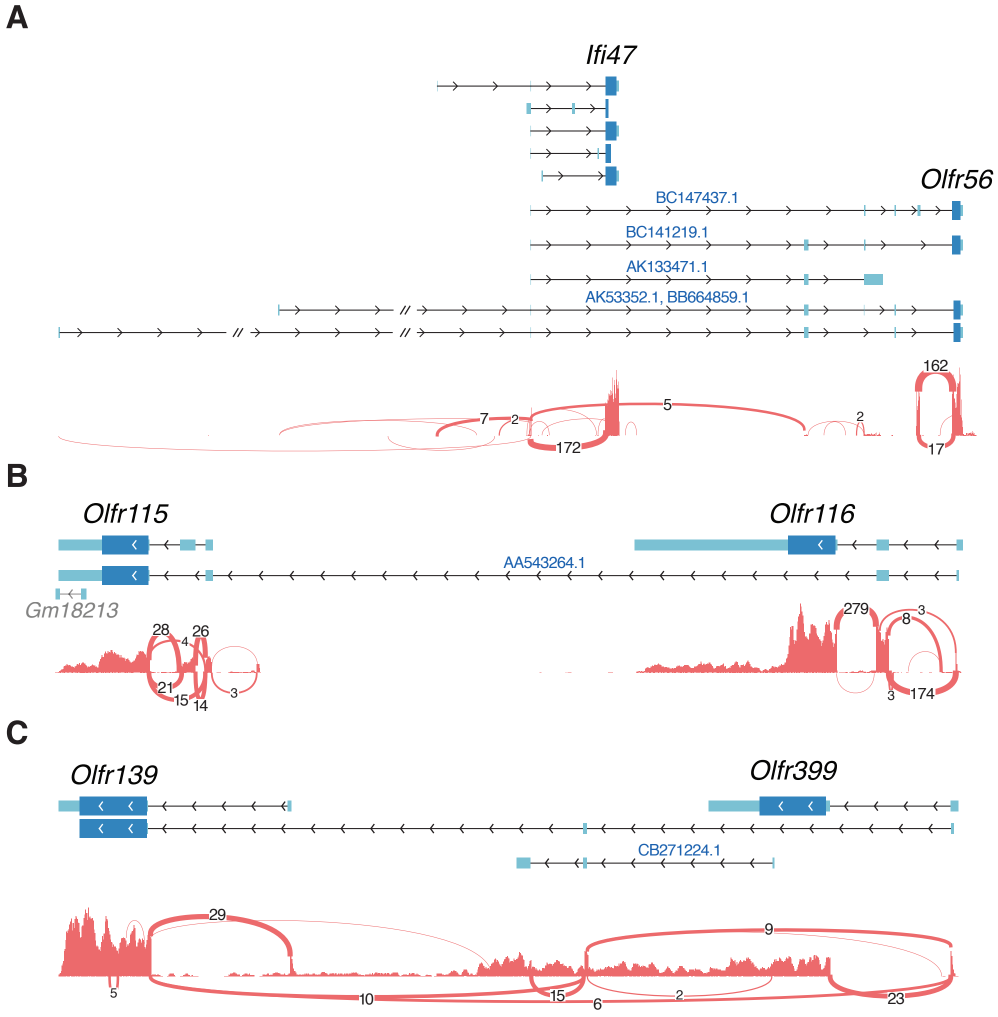


**Supplementary Figure 3 I** Same as Figure 4 but for the additional mouse ORs that share a 5' UTR with a neighbouring gene. mRNA, EST or PacBio clones supporting splice junctions between the two genes are indicated above the corresponding transcript.

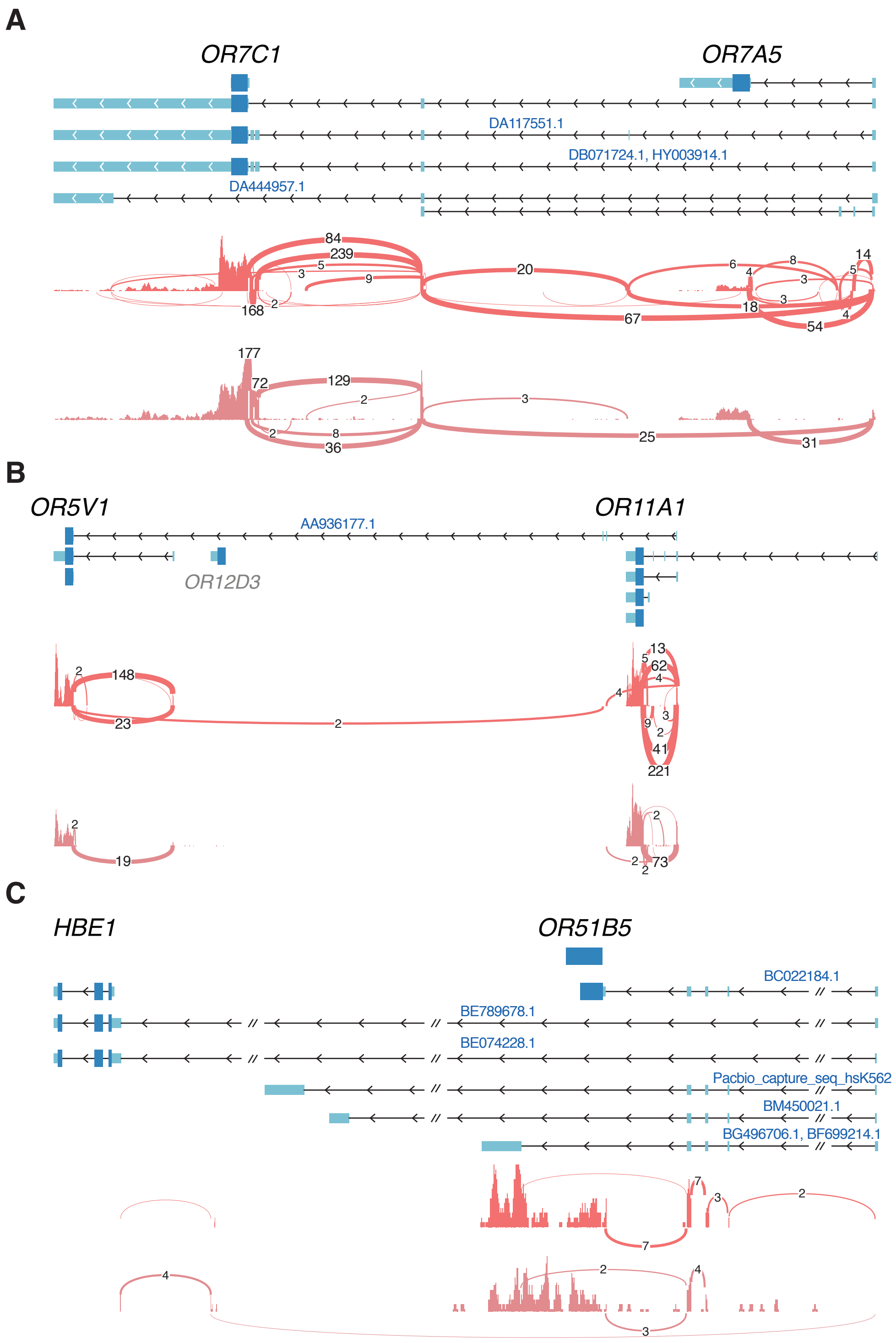


**Supplementary Figure 4 l** Same as Figure 4 but for the additional human ORs that share a 5' UTR with a neighbouring gene. mRNA, EST or PacBio clones supporting splice junctions between the two genes are indicated above the corresponding transcript.

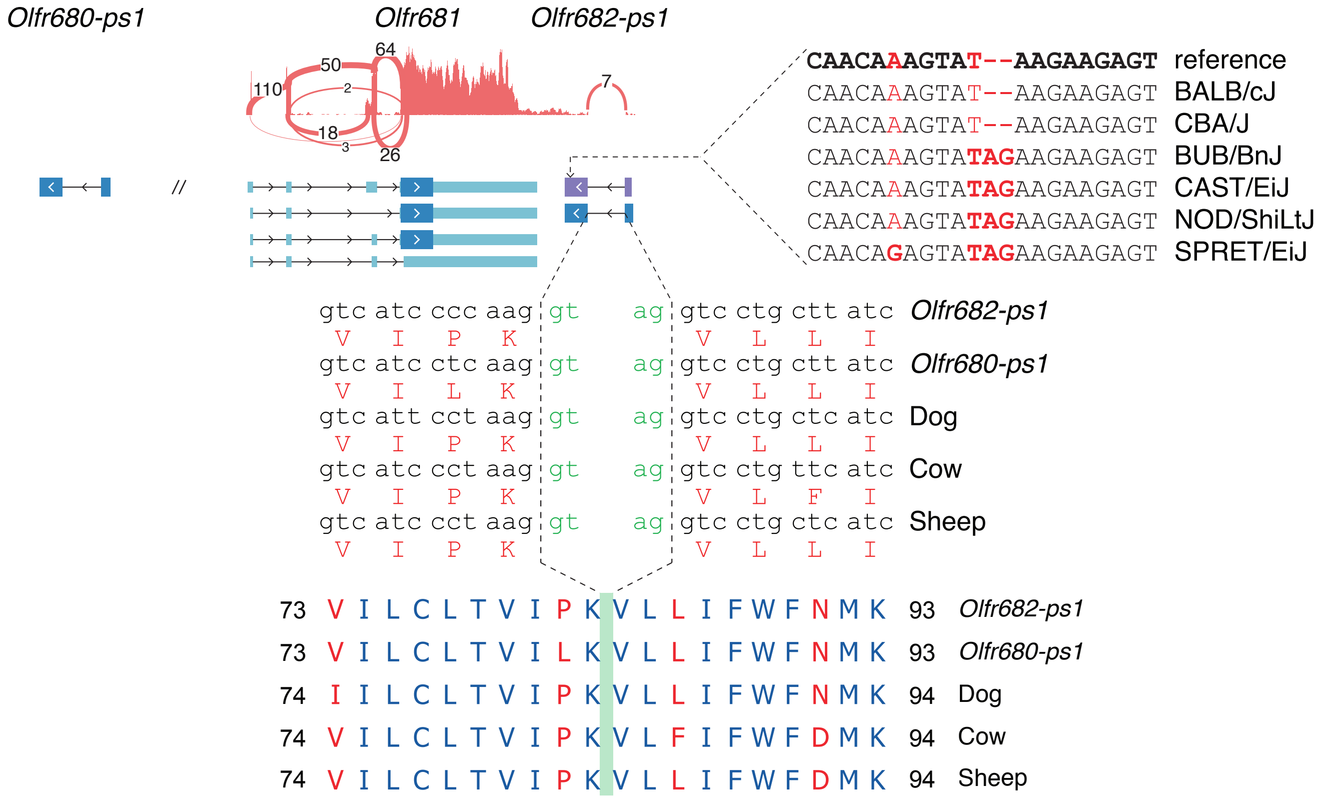


**Supplementary Figure 5 I** Additional example of a split OR gene. On chromosome 7, Olfr682-ps1 was annotated as a pseudogene, but we identified an open reading frame (ORF) spanning two exons that codes for a 311 aa protein. This gene is a polymorphic pseudogene that, in the reference genome, contains a frameshift in the C-terminal domain (purple transcript); however, several mouse strains contain a 2bp indel at position 105,126,541 that restores the correct frame. The splice junction and protein sequence are conserved in several mammals, including dog, cow and sheep. Olfr682-ps1 has a close paralogue, Olfr680-ps1, which shares 97% identity at the protein level. Whereas Olfr680-ps1 lacks transcriptional evidence, we used the conservation with Olfr682-ps1 and other mammals to annotate a full-length split transcript structure.
